# Heterotrimeric G proteins promote FLS2 protein accumulation through inhibition of FLS2 autophagic degradation

**DOI:** 10.1101/438135

**Authors:** Jimi C. Miller, Stacey A. Lawrence, Nicole K. Clay

## Abstract

FLAGELLIN-SENSITIVE 2 (FLS2) is a plant immune receptor that binds bacterial flagellin to activate immune signaling. This immune signal is transduced by a heterotrimeric G protein complex at the plasma membrane and activates downstream signaling. However, it is unknown whether the heterotrimeric G proteins have functions at other subcellular locations away from the plasma membrane. Here, we show that components of the heterotrimeric G protein complex stabilize FLS2 protein levels by inhibiting the autophagic degradation of FLS2. Using genetic analysis, we determined that mutations of G protein components resulted in reduced immune signaling in part due to decreased FLS2 protein levels. Furthermore, reduction of FLS2 protein levels was caused by elevated proteasomal and autophagic degradation of FLS2. Genetic inhibition of autophagy in G protein mutants rescued FLS2 levels and immunity. Our findings suggest that the heterotrimeric G protein components, in addition to being part of the heterotrimeric G protein complex that transduces signals at the plasma membrane, also function away from the plasma membrane to control FLS2 protein levels. These results expand the functional capacity of the heterotrimeric G protein complexes in plant immunity.

## INTRODUCTION

Plants are exposed to a wide range of abiotic and biotic stresses. In response to biotic attack, the plant innate immune system is able to perceive invading pathogens including bacteria and fungi. Plants recognize conserved molecules from pathogens, known as pathogen-associated molecular patterns (PAMPs), such as flagellin from bacteria or chitin from fungal cell walls. PAMPs are perceived at the plasma membrane by pattern recognition receptors (PRRs) that are either receptor-like kinases (RLKs) or receptor-like proteins (RLPs) (Boller and Felix, 2009). Flagellin is recognized by the RLK FLAGELLIN-SENSITIVE 2 (FLS2) receptor and its co-receptor BRASSINOSTEROID INSENSITIVE 1-ASSOCIATED RECEPTOR KINASE 1 (BAK1) (Boller and Felix, 2009; Li et al., 2002). Upon flagellin binding, FLS2 and BAK1 undergo conformational changes and initiate immune signaling. Immune signaling activates a variety of defense responses such as the production of reactive oxygen species (ROS), activation of the mitogen-activated protein kinase (MAPK) cascade, callose deposition, transcriptional reprogramming, and the production of secondary metabolites (Boller and Felix, 2009; Piasecka et al., 2015).

When FLS2 binds bacterial flagellin and activates immune signaling, the plant allocates resources from growth to defense, causing inhibition of plant growth (Boller and Felix, 2009; Gómez-Gómez et al., 2000; Belkhadir et al., 2014). Arabidopsis is able to perceive flagellin at concentrations as low as 3 pM and responds to flagellin in a dose-dependent manner up to a saturating concentration of 1 μM (Felix et al., 1999; Gómez-Gómez et al., 1999). Minimal bacterial populations in the millions are necessary for bacterial survival and infection of plant hosts (Vidaver and Lambrecht, 2004), and empirical evidence indicates that a concentration of 100 nM flagellin is near the physiological levels that plants experience while under pathogen attack (Vidaver and Lambrecht, 2004; Felix et al., 1999; Gómez-Gómez et al., 2000).

Immune signaling from PRRs is transduced by a heterotrimeric G protein complex, which is comprised of a Gα, a Gβ, and a Gγ subunit (Temple and Jones, 2007). The Gα subunit regulates signal activation through a GDP-GTP cycle. The inactive GDP-bound Gα subunit forms a complex with the Gβγ heterodimer, which then binds to the receptor (Oldham and Hamm, 2008). The activated receptor promotes exchange of GDP for GTP, causing the active GTP-bound Gα subunit to dissociate from the receptor complex and the Gβγ heterodimer. The released Gα and Gβγ subunits then activate downstream effectors (Temple and Jones, 2007). The Gβ and Gγ subunits form an obligate heterodimer through a coiled-coil interaction between the N-terminal α-helices of the Gβ and Gγ subunits (Oldham and Hamm, 2008; Temple and Jones, 2007).

The Gβ and Gγ subunits function to activate defense responses upon pathogen attack. ARABIDOPSIS G-PROTEIN BETA SUBUNIT 1 (AGB1) is critical for immunity as the loss-of-function *agb1* mutant exhibits increased susceptibility to the bacterial pathogen *Pseudomonas syringae* and the fungal pathogens *Alternaria brassicicola*, *Botrytis cinerea*, and *Fusarium oxysporum* (Trusov et al., 2006; Trusov et al., 2007; Trusov et al., 2009; Liu et al., 2013). Interestingly, the *agb1* mutant has only a subtle defect in MAPK activation at saturating levels of flagellin (Liu et al., 2013). HETEROTRIMERIC G PROTEIN GAMMA SUBUNIT 1 and 2 (AGG1 and AGG2) are functionally redundant in immunity as the single loss-of-function mutants *agg1* and *agg2* have no defect in immune signaling at saturating levels of flagellin, while the *agg1 agg2* double-mutant has increased susceptibility to *P. syringae*, *A. brassicicola*, *B. cinerea*, and *F. oxysporum* (Trusov et al., 2006; Trusov et al., 2007; Trusov et al., 2009; Liu et al., 2013). Furthermore, the *agb1* mutant and *agg1 agg2* double-mutant have immune signaling defects such as reduction of ROS production and a slight decrease only in MITOGEN-ACTIVATED PROTEIN KINASE 4 (MPK4) activation in response to saturating levels of flagellin (Liu et al., 2013). Because the *agb1* mutant and the *agg1 agg2* double-mutant have the same immunity phenotypes, it appears that the Gβ and Gγ subunits share the same functions in signaling, as expected if they form an obligate heterodimer (Trusov et al., 2006; Trusov et al., 2007; Trusov et al., 2009; Liu et al., 2013).

Some of the environmental stresses that plants experience, such as high temperature, salt stress, and infection by viruses or bacteria, can lead to the accumulation of misfolded or unfolded proteins in the endoplasmic reticulum (ER) (Duwi Fanata et al., 2013; Bao and Howell, 2017). Plants respond to these ER stressors through activation of the unfolded protein response (UPR). The UPR serves to mitigate ER stress by increasing the protein folding capacity of the ER (Liu and Howell, 2010a; Liu and Howell, 2010b; Liu and Howell, 2016; Kørner et al., 2015).

The plant heterotrimeric G proteins are involved in the UPR. The *agb1* and *agg1 agg2* mutants are hypersensitive to tunicamycin, a compound that blocks N-glycosylation and thus induces ER stress (Chen and Brandizzi, 2012; Chakravorty et al., 2015). Moreover, *INOSITOL-REQUIRING PROTEIN 1A* and *1B* (*IRE1A/B)* transcripts are elevated in the *agb1* mutant, suggesting that the Gβγ heterodimer may regulate the UPR (Chen and Brandizzi, 2012). Like the G proteins, the UPR is also involved in immunity. The *ire1a ire1b* double-mutant and the mutant of their downstream target, *BASIC REGION/LEUCINE ZIPPER MOTIF 60* (*bZIP60*), showed increased susceptibility to *P. syringae* expressing the avrRpt2 type III effector (Moreno et al., 2012). This suggests that the UPR, and more specifically *bZIP60*, is an important component of immunity against bacterial pathogens. However, the precise relationship between the G proteins and the UPR in immunity remains unclear (Chen and Brandizzi, 2012).

When misfolded proteins overwhelm the ER protein folding machinery, the UPR activates two cellular protein degradation pathways ER-associated degradation (ERAD) and autophagy (Qi et al., 2017; Liu et al., 2012). ERAD senses misfolded or unfolded proteins in the ER lumen, ubiquinates them, and shuttles them out of the ER to the cytoplasm to be degraded by the proteasome (Liu and Li, 2014; Strasser, 2018). When FLS2 fails to fold properly, it is degraded by ERAD (Lu et al., 2009; Häweker et al., 2010; Liu and Li, 2014). Different mutations in the ectodomain of FLS2 trigger its degradation by ERAD and subsequently the proteasome, causing immune signaling deficiencies (Häweker et al., 2010).

Autophagy is a process that degrades cellular debris accumulated in double-membraned vesicles that are targeted to the vacuole (Marshall and Vierstra, 2018). Autophagy is composed of four major steps: vesicle nucleation; phagophore expansion and closure; delivery of autophagosome to vacuole; and breakdown of the autophagic membrane and degradation of its contents (Marshall and Vierstra, 2018). More than 40 autophagy-related genes (ATG) have been identified that function in autophagy. Autophagy also has the capability to selectively degrade specific cargo, known as selective autophagy. The AUTOPHAGY-RELATED GENE 8 (ATG8) drives cargo selectivity in autophagy by associating with autophagic receptors that are specific for different cargo (Dikic, 2017; Farre and Subramani, 2016; Li and Vierstra, 2012). When autophagy is activated, ATG8 is modified and lipidated to coat the expanding phagophore (Marshall and Vierstra, 2018). Specifically, ATG8 is lipidated through an ubiquitin-like conjugation process that includes ATG7 and ATG3, which are essential for autophagic body formation (Yoshimoto et al., 2004; Suttangkakul et al., 2011; Kim et al., 2013). Unlike ATG8, which is encoded by multiple genes, ATG7 and ATG3 are each encoded by a single gene (Phillips et al., 2008).

Autophagy also plays a role in immunity. Autophagy is essential to activate programmed host cell death (PCD) and restrict PCD only to infected tissue upon pathogen infection (Lenz et al., 2011; Hofius et al., 2009; Liu et al., 2005; Patel et al., 2008; Yoshimoto et al., 2009). Knock down of *ATG6* and *ATG7* genes in tobacco leaves infected with the tobacco mosaic virus (TMV) resulted in PCD that spread beyond infected regions and to uninfected tissue (Patel and Dinesh-Kumar, 2008). Similarly, Arabidopsis plants with reduced *ATG6* expression and the *atg5* loss-of-function mutant had PCD spread beyond the infected regions when infected with the avirulent bacterial pathogen *Pseudomonas syringae* pv. *tomato* (*Pto*) DC3000 carrying the AvrRpm1 effector protein (Yoshimoto et al., 2009; Lenz et al., 2011). The mechanism by which autophagy affects immunity remains unknown, but autophagy appears essential to ensure an appropriate defense response.

In this study, we show that the reduction of immune signaling in the *agg1 agg2* double-mutant is in part due to decreased FLS2 protein levels. We found that the *agg1 agg2* mutant had increased FLS2 protein degradation, while *FLS2* transcripts were unaffected. Simultaneous chemical inhibition of both the proteasome and autophagy was sufficient to rescue levels of functional FLS2 and growth inhibition by flagellin. Furthermore, disabling autophagy alone (without disabling the proteasome) in the *agg1 agg2* mutant was sufficient to restore FLS2 and immune signaling back to wild-type levels. Interestingly, AGG1 and AGG2 localized at the ER membrane rather than at autophagic bodies. Our results indicate that the Gβγ heterodimer promotes FLS2 levels by inhibiting autophagic degradation of FLS2 at the ER membrane, in contrast to expectations that the heterotrimeric G proteins only function at the plasma membrane.

## RESULTS

### Plant immune signaling is impaired in the *agg1 agg2* mutant at physiological levels of flagellin

Previous reports have shown that the Gβ single mutant, *agb1*, and the Gγ double-mutant, *agg1 agg2,* show subtle defects in immune signaling at saturating concentrations of flagellin (Trusov et al., 2007; Liu et al., 2013). However, the extent of immune signaling reduction in these G protein mutants may differ at physiological concentrations of flagellin. To test if the heterotrimeric G proteins are crucial in immune signaling at physiological concentrations of flagellin, we measured growth inhibition upon flagellin (flg22) treatment in wild type and G protein mutants. We treated seedlings with 100 nM flg22, a concentration near physiological levels, and compared the mass to that of mock-treated seedlings. Treatment with 100 nM flg22 caused similar growth inhibition in both the *agb1* mutant and in the wild type, indicating that the perception of flagellin at physiological levels is not impaired in the *agb1* mutant (Figure S1A). However, the *agg1 agg2* double mutant did show reduced growth inhibition (20% growth inhibition) upon treatment with 100 nM flg22, which is intermediate between wild type (55% growth inhibition) and a T-DNA insert mutant in the promoter of *FLS2* that severely reduces *FLS2* transcript levels (Zipfel et al., 2004), *fls2c* (noted here afterward as *fls2*) (6% growth inhibition) (Figure 1A). This result suggests that Gγ subunits play an important role in immune signaling at near-physiological concentrations of flagellin.

**Figure 1.**
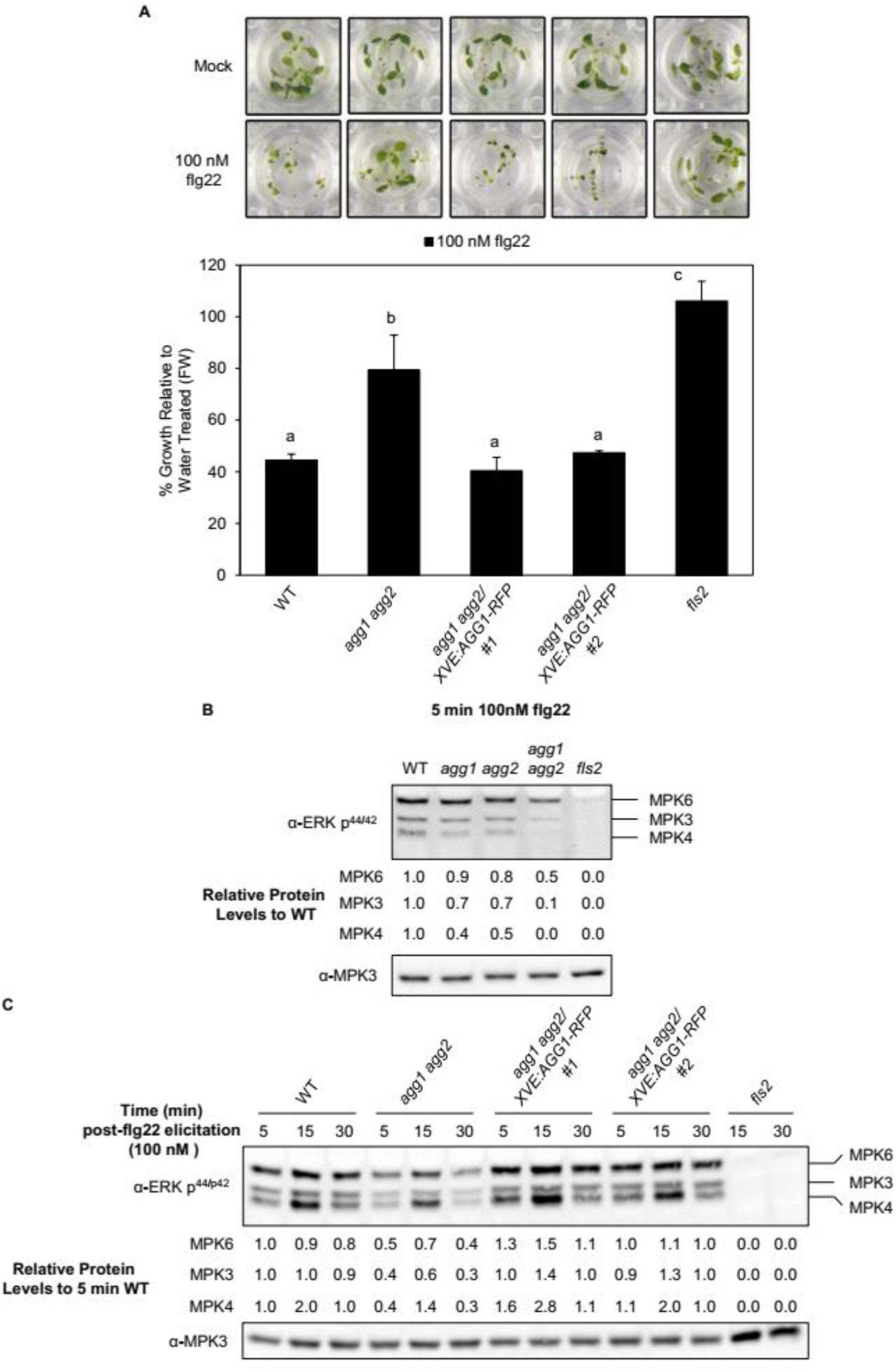
Plant defense signaling is impaired in the *agg1 agg2* mutant. (A) Growth inhibition analysis of 9-day-old seedlings pre-treated with 20 μM β-estradiol and water (control) or 20 μM β-estradiol plus 100 nM flg22 for 6 days. Data represent mean ± SD of four (*fls2*) or five (all others) replicates of five seedlings. (B) Immunoblot analysis of phosphorylated, active MAPKs in 9-day-old WT*, agg1, agg2, agg1 agg2, fls2* in response to 100 nM flg22 for 5 minutes. (C) WT*, agg1 agg2, agg1 agg2/XVE:AGG1-RFP* lines, and *fls2* in response to 100 nM flg22 for 5, 15, and 30 minutes. (B-C) Numbers under immunoblots indicate phosphorylated MPK3, MPK4, and MPK6 signal intensities normalized to those of total MPK3 (loading control) and then normalized to WT 5 minutes post-elicitation.

Previous studies using saturating concentrations of flagellin showed that MAPK activation was only slightly reduced in mutants defective in the Gβ or Gγ subunit genes (Liu et al., 2013). However, it is unclear whether the heterotrimeric G proteins are necessary to activate the MAPK cascade at physiological concentrations of flagellin. We tested MAPK activation at near-physiological flagellin concentrations by measuring the amount of phosphorylated MPK6, MPK3, and MPK4 in the wild type and the G protein mutants. In accordance with the previous data (Liu et al., 2013), the *agb1* mutant only had decreased MPK4 activation upon physiological flg22 treatment, while MPK6 and MPK3 were unaffected (Figure S1B). The previous report also showed that only MPK4 activation was decreased in *agg1 agg2* at saturating flg22 concentrations, while MPK6 and MPK3 were unaffected (Liu et al., 2013). However, at near-physiological flagellin concentrations, the *agg1 agg2* mutant showed decreased MPK6, MPK3, and MPK4 activation compared to wild type, but not fully ablated as in the *fls2* mutant (Figure 1B). MAPK activation at near-physiological flagellin concentrations was slightly reduced in the *agg1* and *agg2* single mutants compared to wild type (Figure 1B). This is also differs from the previous report of which MAPK activation was unaffected in the *agg1* and *agg2* single mutants at saturating flg22 concentrations (Liu et al., 2013). Together, this result suggests that the Gγ subunits, AGG1 and AGG2, are critical for the activation of MAPK at near-physiological levels of flagellin.

Callose is a glucan polymer that is deposited at the site of pathogen contact to prevent pathogen entry (Boller and Felix, 2009). Callose deposition is a MAPK-dependent, long-term immune response, which can be used as a quantitative physiological marker of responses to flg22 treatment (Frei dit Frey et al., 2014; Clay et al., 2009). To test if the heterotrimeric G proteins are required for callose deposition at near-physiological concentrations of flg22, we quantified the number of callose deposits in wild type and the G protein mutants in 9-day-old seedlings after flg22 elicitation. Both the *agb1* and the *agg1 agg2* mutants had significantly fewer number of callose deposits compared to wild type (Figure S1C). However, the *agg1 agg2* double-mutant had a more severe phenotype than the *agb1* single mutant. Together, this suggests that the heterotrimeric G proteins are necessary for proper callose deposition at near-physiological concentrations of flg22.

The *agb1* mutant exhibited a weaker phenotype in our assays when compared to the *agg1 agg2* double-mutant, suggesting that there may be additional redundant Gβ subunits (Miller et al., 2019). Therefore, we focused our work on the Gγ subunits AGG1 and AGG2 which showed consistent phenotypes. To verify that the phenotypes in the *agg1 agg2* mutant were caused by the loss of AGG1 and AGG2, we performed a rescue experiment by creating a transgenic line with inducible *AGG1* expression. The transgenic line includes *AGG1-RFP* under a β-estradiol-inducible promoter expressed in the *agg1 agg2* background. Using this line, we induced *AGG1-RFP* expression and measured seedling growth inhibition, MAPK activation, and callose deposition in the presence or absence of 100 nM flg22. We were able to restore flg22 growth inhibition back to wild-type levels after induction of *AGG1-RFP* in two independent transgenic lines (Figure 1A). Expression of *AGG1-RFP* also rescued MAPK activation back to wild-type levels (Figure 1C). Interestingly, these transgenic lines were able to increase the number of callose deposits upon flg22 treatment but not back to wild-type levels, suggesting that expression of *AGG1-RFP* is unable to fully rescue all of the *agg1 agg2* mutant phenotypes (Figure S1C). These data, in combination with the above experiments, suggest that at physiological levels of flagellin the heterotrimeric G proteins are required to activate immune signaling.

### Total FLS2 protein levels are reduced in the *agg1 agg2* mutant

Induction of *FLS2* transcripts upon bacterial pathogen infection requires the heterotrimeric G proteins, whereas FLS2 protein accumulation under normal growth conditions is unaffected by the loss of AGB1 (Liu et al., 2013; Lee et al., 2013). In order to investigate how the Gγ subunits affect immune signaling, we measured total FLS2 protein levels under normal growth conditions in wild type, the *agg1 agg2* double-mutant, and our *agg1 agg2*/*AGG1-RFP* rescue transgenic seedlings. Interestingly, total FLS2 protein levels were reduced in the *agg1 agg2* mutant, indicating that both Gγ subunits, AGG1 and AGG2, affect FLS2 protein levels. Induction of *AGG1-RFP* expression in the *agg1 agg2* mutant fully restored FLS2 protein levels (Figure 2). This result shows that the Gγ subunits AGG1 and AGG2 affect FLS2 protein accumulation, suggesting that lowered FLS2 levels contributed to the lowered immune signaling.

**Figure 2.**
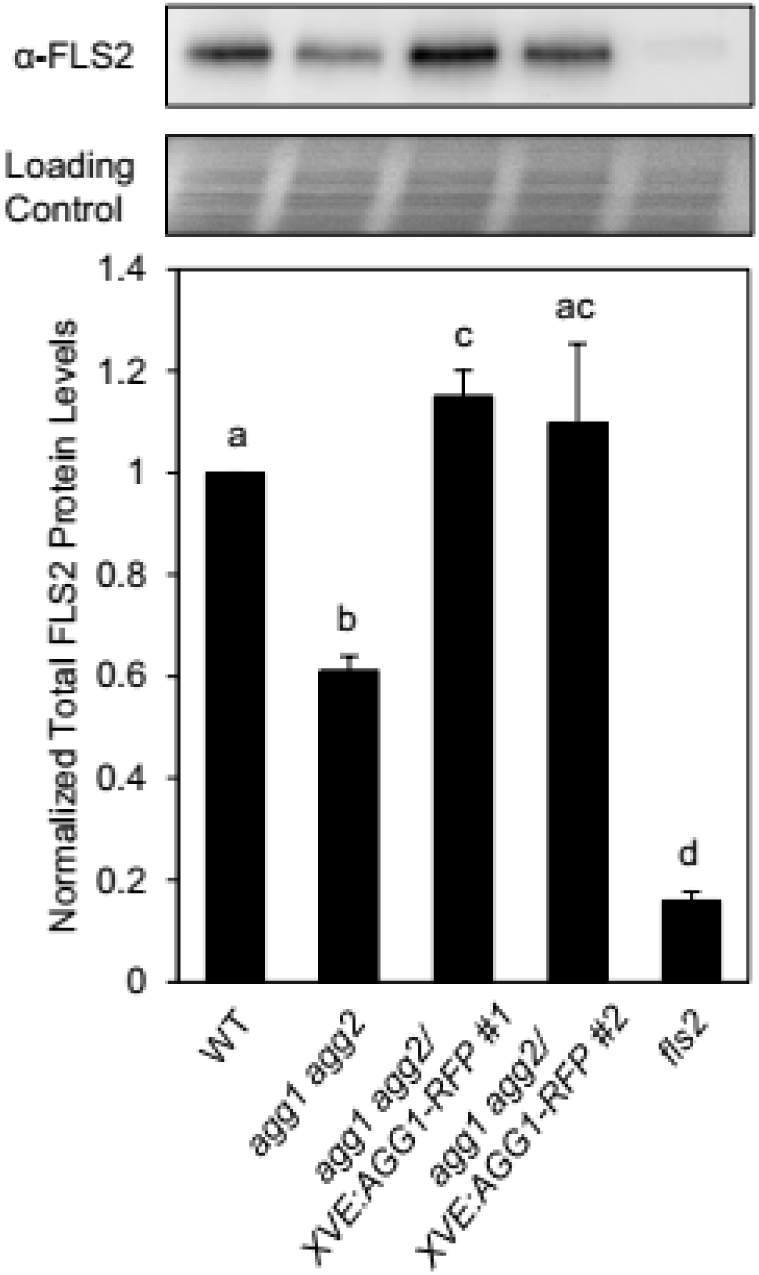
Total FLS2 protein levels are reduced in the *agg1 agg2* mutant. Immunoblot analysis of FLS2 protein in 9-day-old seedlings treated with β-estradiol. Data in graph represent the mean ± SD of three replicates of twelve seedlings. Letters in graph indicate significant differences from WT (*P*-value <0.05, two-tailed *t* test).

### AGG1 and AGG2 modulate total FLS2 protein levels post-transcriptionally

Since loss of AGG1 and AGG2 causes a reduction in total FLS2 protein levels, we next wanted to know if this effect is transcriptional or post-transcriptional. We performed a comprehensive transcriptional and protein analysis of FLS2 in wild type and *agg1 agg2*. We collected tissue from the wild-type and *agg1 agg2* mutant across a 24 hour time course (12 hour light/ 12 hour dark) and performed qRT-PCR and western blot to track the levels of *FLS2* mRNA and protein. *FLS2* transcript levels in the *agg1 agg2* mutant did not differ greatly from wild type, although at some time points there were small significant in changes transcripts (Figure 3A). In contrast, the levels of the FLS2 protein were about two-fold lower at dawn in the *agg1 agg2* mutant compared to those of wild type (Figure 3B). These results suggest that the reduction of FLS2 protein levels in the *agg1 agg2* mutant occurs through a post-transcriptional mechanism.

**Figure 3.**
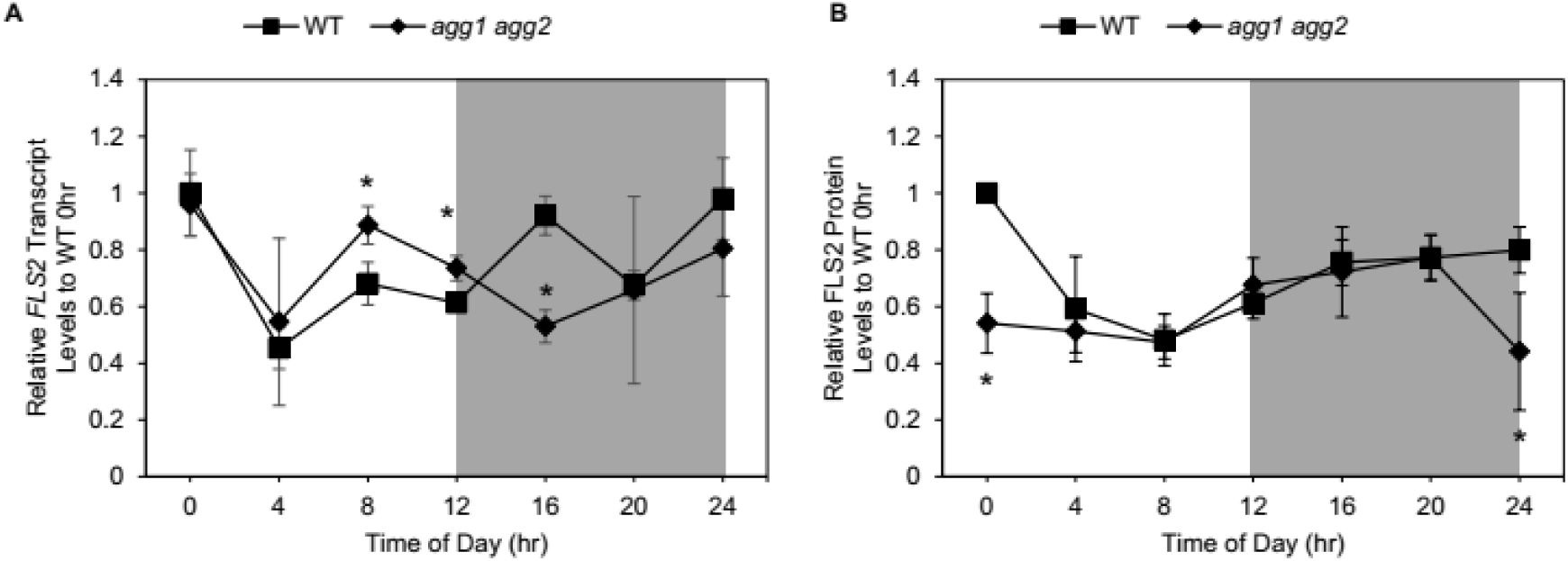
AGG1 and AGG2 modulate total FLS2 protein levels post-transcriptionally. (A) Time course of *FLS2* transcript levels in WT and *agg1 agg2* 9-day-old seedlings. Data represent mean ± SD of four replicates of twelve seedlings. Transcript levels were normalized to WT at 0hr. Asterisks indicate significant differences from WT (*P*-value <0.05, two-tailed *t* test). Shading indicates night. (B) Time course of FLS2 protein levels in WT and *agg1 agg2* 9-day-old seedlings. Data represent the mean ± SD of three replicates of twelve seedlings. Protein levels were normalized to WT at 0hr. Asterisks indicate significant differences from WT. (*P*-value <0.05, two-tailed *t* test). Shading indicates night.

### AGG1 and AGG2 regulate FLS2 protein accumulation by regulating FLS2 protein degradation

AGG1 and AGG2 regulate total FLS2 levels post-transcriptionally, however, it remains unknown if this regulation occurs post-translationally. To determine whether AGG1 and AGG2 affect FLS2 levels post-translationally, we wanted to investigate the stability of FLS2 protein in the *agg1 agg2* mutant. We treated seedlings with cycloheximide to block protein synthesis and then measured FLS2 protein levels at 0, 8 and 16 hours after cycloheximide treatment. At 8 and 16 hours post-cycloheximide treatment, FLS2 protein levels in the *agg1 agg2* mutant were considerably reduced compared to wild type (Figure 4A). This suggests that AGG1 and AGG2 affect the degradation of FLS2 protein.

**Figure 4.**
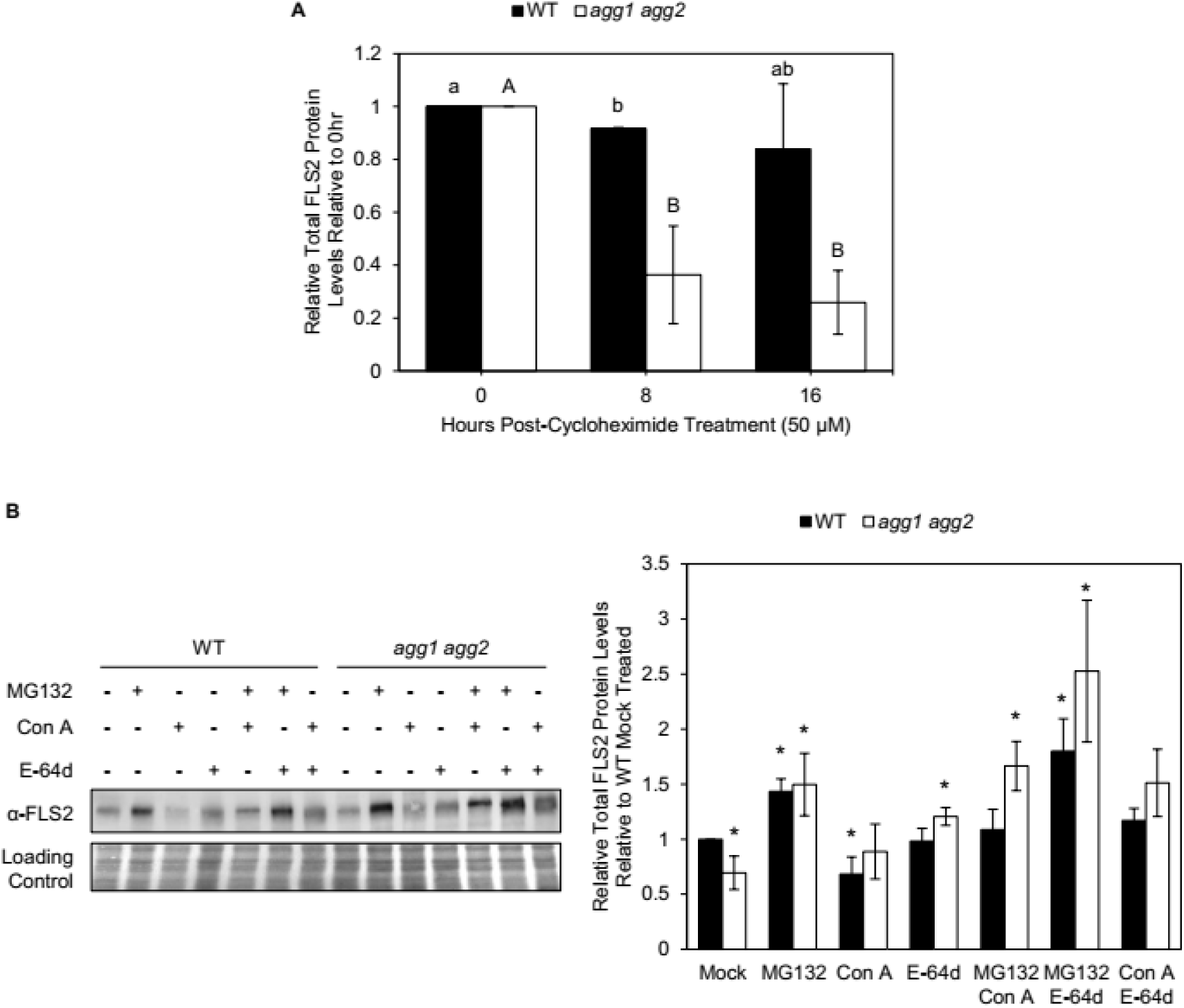
AGG1 and AGG2 affect FLS2 protein degradation. (A) Time course of FLS2 protein levels in WT and *agg1 agg2* 9-day-old seedlings in response to 50 µM cycloheximide. Data represent mean ± SD of three replicates of twelve seedlings. Different letters in indicate significant differences (*P*-value <0.05, two-tailed *t* test). (B) Immunoblot analysis of FLS2 protein in WT and *agg1 agg2* 9-day-old seedlings treated with DMSO, 50 µM MG132, 20 µM E-64d, and/or 2 µM Concanamycin A (Con A) for 24 hr. Data represent mean ± SD of three replicates of twelve seedlings. Asterisks in graph indicate significant differences from mock-treated WT (*P*-value <0.05, two-tailed *t* test).

FLS2 is degraded by the proteasome (Lu et al., 2011), and the results above suggest that the Gγ subunits are able to affect FLS2 protein accumulation by inhibiting the protein degradation of FLS2. However, it remains unknown if the Gγ subunits inhibit the degradation of FLS2 via the proteasome pathway. To determine if the Gγ subunits inhibit the proteasomal degradation of FLS2, we measured FLS2 protein levels in seedlings treated with the proteasome inhibitor MG132. If FLS2 is being degraded by the proteosome to a greater extent in the *agg1 agg2* mutant than in wild type, then MG132 treatment in *agg1 agg2* should restore FLS2 protein levels. In our mock-treatment, FLS2 levels were lower in *agg1 agg2* compared to wild type. MG132 treatment in wild-type seedlings increased FLS2 levels beyond those in the mock-treated wild type (Figure 4B). This is due to the basal level of FLS2 recycling and degradation that is needed to maintain proper FLS2 levels at the plasma membrane and prevent inadvertent auto-activation of FLS2 in the absence of flagellin (Robatzek et al., 2006; Beck et al., 2012). MG132 treatment in the *agg1 agg2* mutant rescued FLS2 protein levels beyond those in the mock-treated wild type. FLS2 levels in the *agg1 agg2* mutant were similar to those in the wild type treated with MG132 (Figure 4B). Our result with MG132 suggests that the Gγ subunits affect FLS2 levels by inhibiting the proteasomal degradation of FLS2.

FLS2 is also recycled through multivesicular bodies and targeted by autophagy (Robatzek et al., 2006; Beck et al., 2012; Yang et al., 2018). However, it remains unknown if the Gγ subunits inhibit the degradation of FLS2 via autophagy. To determine if the Gγ subunits inhibit the autophagic degradation of FLS2, we measured FLS2 protein levels in seedlings treated with the autophagy inhibitors concanamycin A (Con A), E64-d, or both inhibitors. In Con A-treated wild-type seedlings, FLS2 levels were lower than the mock-treated wild type, which may be due to the toxicity of Con A at these levels. FLS2 levels in the *agg1 agg2* mutant treated with Con A were similar to those in the mock-treated wild type (Figure 4B). In E64-d treated seedlings, FLS2 levels in wild type were similar to the mock-treated wild type, whereas FLS2 levels in the *agg1 agg2* mutant after E64-d treatment were increased beyond the mock-treated wild type (Figure 4B). Furthermore, when seedlings were treated with both Con A and E64-d, FLS2 levels in both wild type and *agg1 agg2* were elevated compared to those in the mock-treated wild type (Figure 4B). These Con A and E64-d results suggest that the Gγ subunits affect FLS2 levels by inhibiting the autophagic degradation of FLS2. Our results above indicate that the Gγ subunits inhibit the protein degradation of FLS2 via either the proteasome or autophagy. In order to further investigate how the Gγ subunits inhibit degradation of FLS2, we treated seedlings with both MG132 and Con A or MG132 and E64-d. Wild-type seedlings treated with MG132 and Con A had FLS2 levels similar to those in the mock-treated wild type. Treatment of MG132 and Con A in *agg1 agg2* seedlings increased FLS2 levels beyond those in the mock-treated wild type (Figure 4B). Wild-type seedlings treated with MG132 and E64-d had FLS2 levels similar to those in the mock-treated wild type. Treatment of MG132 and E64-d in *agg1 agg2* seedlings increased FLS2 levels beyond those in the mock-treated wild type. Interestingly, *agg1 agg2* seedlings treated with MG132 and E64-d increased FLS2 levels beyond any other treatment tested (Figure 4B). Taken together, our results suggest that the Gγ subunits are necessary to inhibit both the proteasomal and autophagic degradation of FLS2 and to thus maintain adequate FLS2 levels.

### Inhibiting the proteasome and autophagy in *agg1 agg2* rescues seedling growth inhibition upon flg22 elicitation

Inhibiting the proteasome and autophagy in the *agg1 agg2* mutant was sufficient to restore FLS2 protein levels. However, it remains unclear if the restored FLS2 protein is able to recognize flagellin and rescue the *agg1 agg2* mutant phenotype. To test the functionality of the restored FLS2 in the *agg1 agg2* mutant, we quantified flagellin growth inhibition by measuring the mass of seedlings treated with or without flagellin and in the absence or presence of MG132, E-64d, or both. However, due to the toxic effects these inhibitors may have on the plants, we optimized the concentrations of both MG132 and E-64d to ensure they did not affect the growth of the wild-type seedlings (Figure 5A). Next, we applied the inhibitors at these optimized concentrations to wild type and the *agg1 agg2* mutant seedlings in the presence or absence of flg22. Treatment with either MG132 or E64-d alone was unable to restore the ability of the *agg1 agg2* mutant to inhibit growth upon flg22 treatment to wild-type levels. Treatment with both MG132 and E64-d was sufficient to restore growth inhibition upon flg22 elicitation in the *agg1 agg2* mutant to those seen in the wild type (Figure 5B). This result indicates that AGG1 and AGG2 prevent the proteasomal and autophagic degradation of FLS2, and the reduction of FLS2 levels in the *agg1 agg2* mutant partially contributes to the immune signaling defects in response to flagellin seen in our experiments.

**Figure 5.**
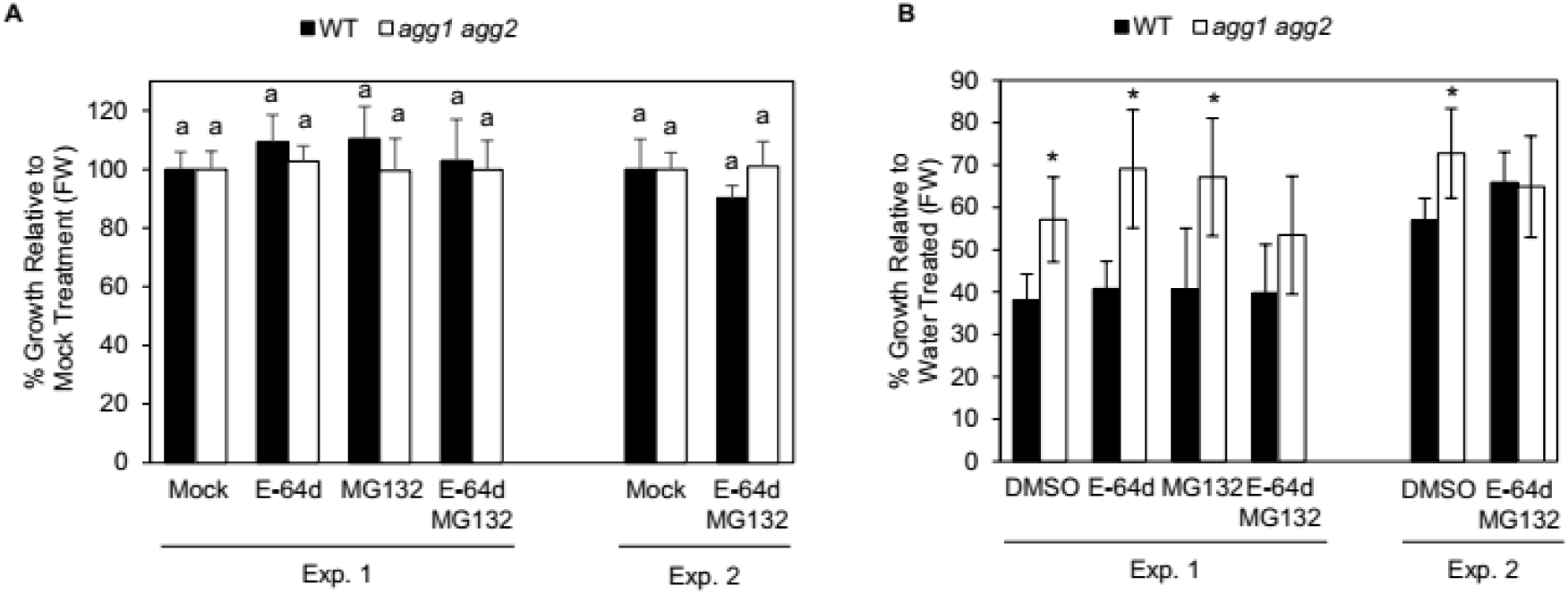
Inhibiting proteasome and autophagy in *agg1 agg2* rescues seedling growth inhibition upon flg22 elicitation. (A) Two independent growth analyses of 9-day-old plants treated with water and with DMSO (mock), 50 nM MG132, 20 nM E-64d, or 50 nM MG132 and 20 nM E64-d for 4 days. Data represent mean ± SD of five replicates of five seedlings. Different letters indicate significant differences from WT (*P*-value <0.05, two-tailed *t* test). (B) Two independent growth inhibition analyses of WT and *agg1 agg2* 9-day-old seedlings treated with water (control) or 100 nM flg22 for two days and then treated with DMSO (mock), 50 nM MG132, 20 nM E-64d, or 50 nM MG132 and 20 nM E64-d for 4 days. Data represent mean ± SD of five replicates of five seedlings. Asterisks indicate significant differences from WT (*P*-value <0.05, two-tailed *t* test).

### Genetic inhibition of autophagy confirms that AGG1 and AGG2 regulate FLS2 degradation via autophagy

Our results have shown that chemical inhibition of the proteasome and autophagy degradation pathways in the *agg1 agg2* mutant is sufficient to rescue functional FLS2 protein levels. However, it remains unclear whether inhibiting the proteasome or autophagy genetically in the *agg1 agg2* background is able to phenocopy the results observed in our chemical inhibition experiments. To test this, we wanted to cross the *agg1 agg2* double-mutant with proteasome or autophagy mutants. However, we were unable to obtain mutants of core proteasomal mutants, and were restricted to testing autophagy mutants in a genetic cross into the *agg1 agg2* mutant. To ensure that the autophagic machinery is functionally ablated, we chose the *atg7* and *atg3* single mutants to cross into the *agg1 agg2* background as they are essential for autophagy (Kim et al., 2013; Suttangkakul et al., 2011). To show that flagellin perception in either the *agg1 agg2 atg7* or *agg1 agg2 atg3* triple mutants were restored back to wild-type levels, we measured the growth inhibition in these mutants upon flg22 treatment. Growth was inhibited approximately 10% more in the *agg1 agg2 atg7* and *agg1 agg2 atg3* mutants (56% and 59%, respectively) than in the wild type (45%) upon flagellin treatment (Figure 6A). This data suggests that *AGG1* and *AGG2* genetically interact with autophagy to inhibit the autophagic degradation of FLS2.

**Figure 6.**
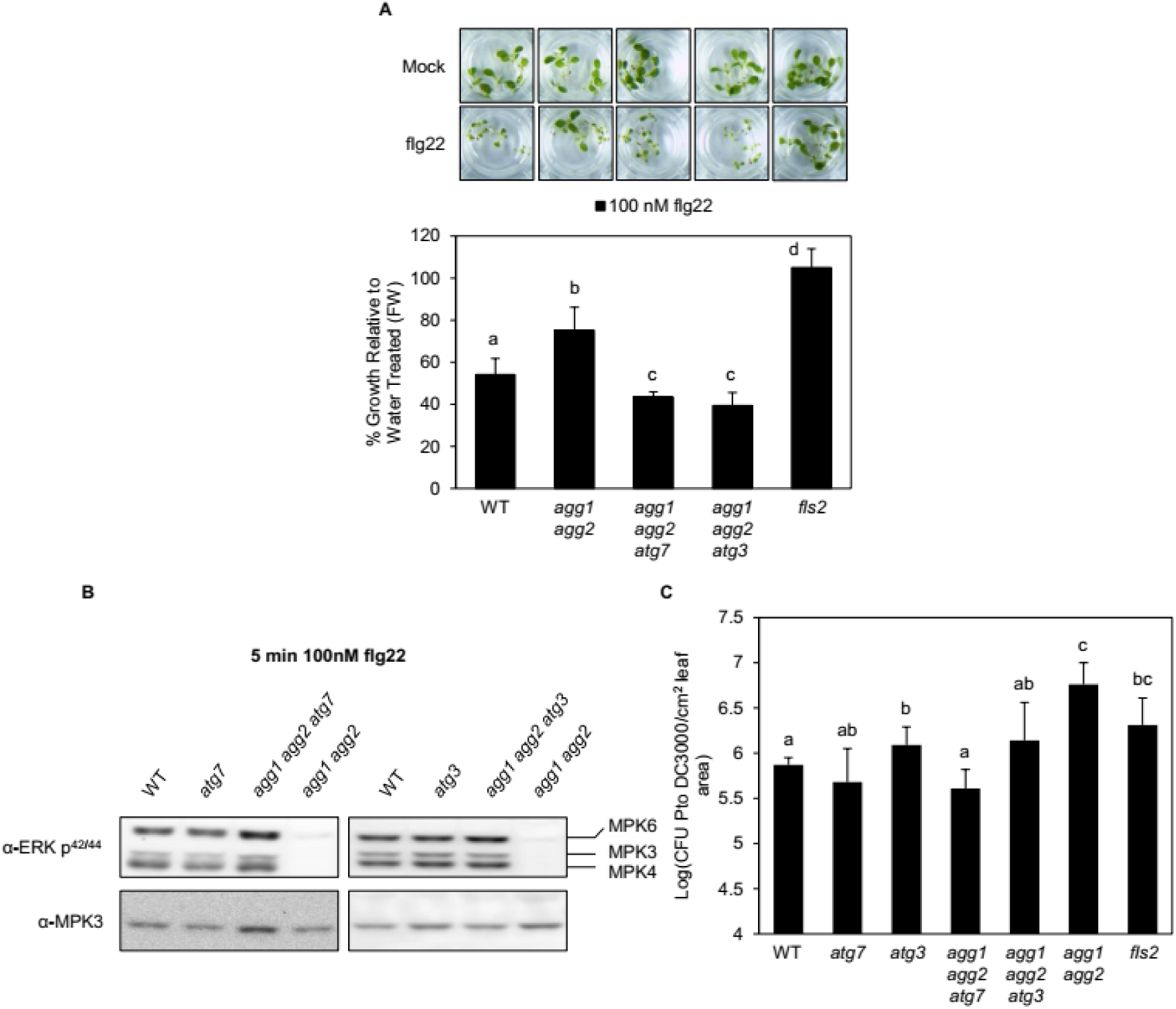
Knockout of autophagy proteins ATG7 or ATG3 in the *agg1 agg2* mutant rescues *agg1 agg2* phenotype. (A) Growth inhibition analysis of 9-day-old seedlings treated with water (control) or 100 nM flg22 for 6 days. Data represent mean ± SD of four (*fls2*) or five (all others) replicates of five seedlings. (B) Immunoblot analysis of phosphorylated, active MAPKs in 9-day-old WT*, atg7, atg3, agg1 agg2, agg1 agg2 atg7,* and *agg1 agg2 atg3* in response to 100 nM flg22 for 5 minutes. (C) Growth analysis of bacterial pathogen *Pto* DC3000 in 5-week-old surface-inoculated leaves. Data represent mean ± SD of six replicates. Different letters indicate significant differences (*P*-value <0.05, two-tailed *t* test).

While flagellin growth inhibition is restored in the *agg1 agg2 atg7* and *agg1 agg2 atg3* triple mutants, it remains unknown whether downstream immune signaling is also restored in these mutants. We tested downstream MAPK activation in these triple mutants upon flg22 treatment. We measured the amount of phosphorylated MPK6, MPK3, and MPK4 in seedlings after elicitation with 100 nM flg22. MAPK activation was restored in the *agg1 agg2 atg7* and *agg1 agg2 atg3* mutants back to wild-type levels when treated with flg22. Moreover, in the *atg7* and *atg3* single mutants had no defect in MAPK activation upon flg22 treatment (Figure 6B). This data suggests that AGG1 and AGG2 genetically interact with the autophagy machinery to inhibit the autophagic degradation of FLS2 and promote immune signaling in response to physiological levels of flg22.

The *agg1 agg2* double mutant is highly susceptible to the bacterial pathogen *Pseudomonas syringae Pto* DC3000 (Liu et al., 2013). It remains unknown whether the *agg1 agg2 atg7* and *agg1 agg2 atg3* triple mutants are able to rescue the susceptibility phenotype of the *agg1 agg2* double-mutant back to wild type. To determine if the *agg1 agg2 atg7* and *agg1 agg2 atg3* triple mutants rescue the *agg1 agg2* susceptibility phenotype, we infiltrated adult leaves with *P. syringae* DC3000 and quantified the number of bacteria in the leaf. The *atg7* mutant had a similar phenotype to wild type, whereas the *atg3* mutant was slightly more susceptible than wild type to *Pto* DC3000 (Figure 6C). The *agg1 agg2 atg7* and *agg1 agg2 atg3* triple mutants were able to rescue the *agg1 agg2* susceptibility phenotype against *Pto* DC3000 back to wild-type levels (Figure 6C). This suggests that disrupting autophagy in the *agg1 agg2* mutant restores functional FLS2 levels and in turn restores immune signaling back to wild-type levels. Together, these results suggest that *AGG1* and *AGG2* genetically interact with the autophagy pathway to prevent the degradation of FLS2, resulting in stable FLS2 protein levels and a robust immune response upon pathogen attack.

### AGG1 and AGG2 function and localize to the endoplasmic reticulum membrane

Our results have shown that the Gγ subunits inhibit the autophagic degradation of FLS2. The sub-cellular location where AGG1 and AGG2 inhibit the autophagic degradation of FLS2 remains unknown. To determine the sub-cellular localization of AGG1 and AGG2, we expressed *AGG1* with a C-terminal fluorescent tag under a β-estradiol-inducible promoter in tobacco leaves and imaged their sub-cellular localization using confocal microscopy. AGG1 localized at the ER membrane with the ER protein marker HDEL (Gomord et al., 1997). Moreover, AGG1 co-localized with AGB1 and FLS2 at the ER membrane, suggesting that AGG1 and AGG2 localize and function at the ER membrane, away from the plasma membrane (Figure 7B). However, we never observed AGG1 localizing at puncta in the vacuole, which are indicative of autophagic bodies (Kirisako et al., 1999; Yoshimoto et al., 2004; Contento et al., 2005). We also ensured that the C-terminal tag did not affect the localization of AGG1 by inducing plasmolysis in the tobacco cells with mannitol. This shrinks the vacuole, pulling the plasma membrane away from the cell wall. As the plasma membrane pulls away from the cell wall, sections of the plasma membrane are stuck to the cell wall which create strands called Hechtian strands (Buer et al., 2000). AGG1-GFP co-localized with FLS2-RFP at these Hechtian strands, which indicates that the C-terminal tag did not affect the localization of AGG1 (Figure S2). This result suggests that the Gγ subunits localize to the ER membrane in addition to localizing at the plasma membrane.

**Figure 7.**
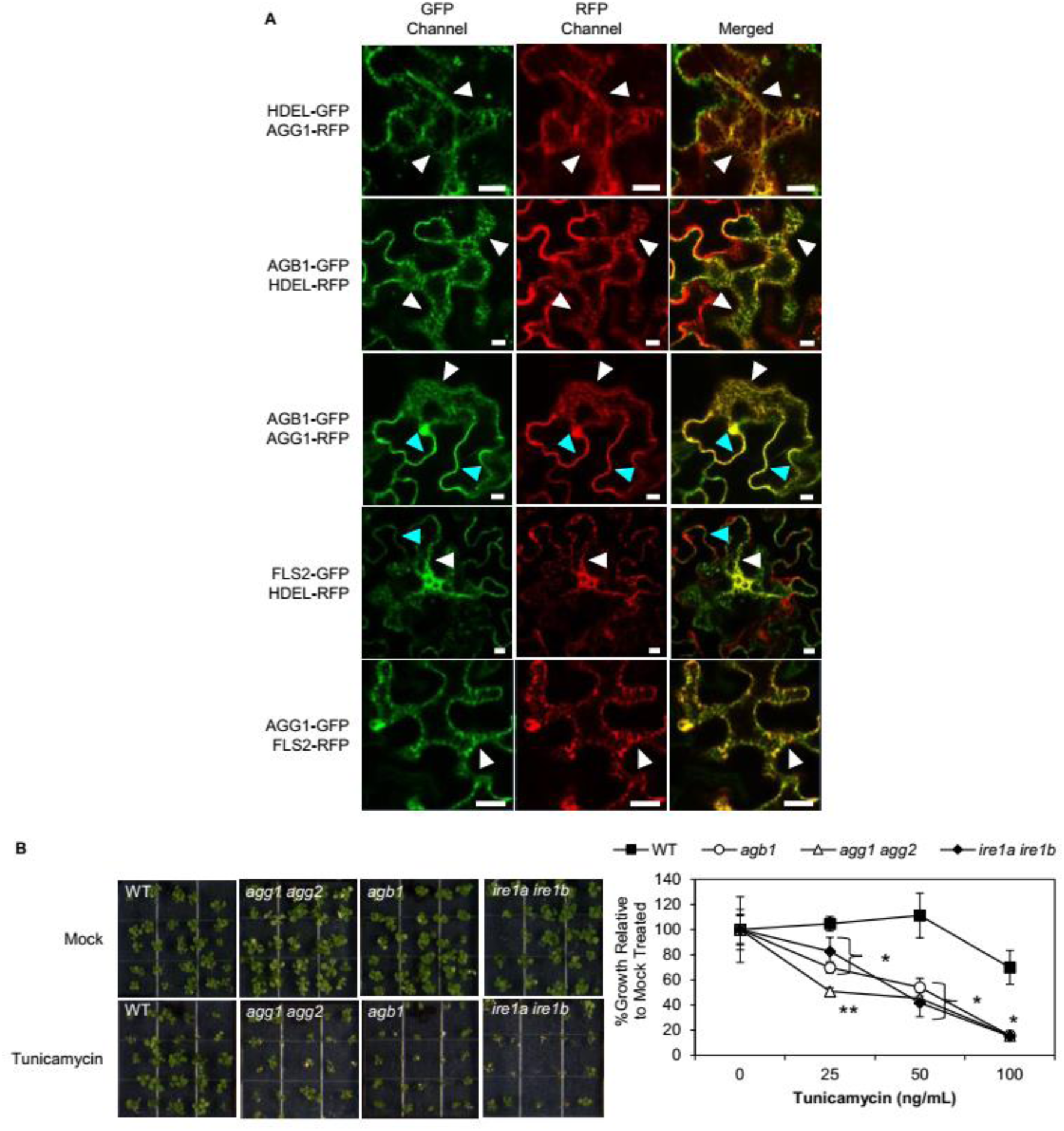
AGG1 and AGG2 localize and function to the endoplasmic reticulum membrane. (A) Co-localization of AGG1 with AGB1 and FLS2 at the ER membrane in transiently transfected *N. benthamiana* leaves pretreated with 20 μM β-estradiol for 4-8 hr. HDEL is an ER marker. White arrow heads indicate ER localization. Light blue arrow heads indicate plasma membrane localization. White bars represent 20 μm. (B) Growth inhibition analysis of 14 day-old plants in response to 0 (control), 25, 50 and 100 ng mL^−1^ tunicamycin. Data represent mean ± SD of five replicates of five seedlings. Asterisks indicate significant differences from WT; double asterisks indicate significant differences from WT, *agb1* and *ire1a ire1b*.

The Gβ and Gγ subunits are involved in the UPR under ER stress at saturating levels of tunicamycin, a compound that causes ER stress and triggers the UPR (Chakravorty et al., 2015). However, the extent of sensitivity in the *agg1 agg2* mutant to ER stress is unknown. To determine the sensitivity of the *agg1 agg2* mutant to tunicamycin treatment, we measured the mass of seedlings grown in the presence of varying concentrations of tunicamycin and compared them to seedlings grown without tunicamycin. Similar to previous reports, the *agg1 agg2* mutant was hypersensitive to tunicamycin (Chakravorty et al., 2015). Interestingly, the *agg1 agg2* mutant was more sensitive to tunicamycin than the *agb1* mutant and the UPR sensor double-mutant *ire1a ire1b*, which are sensitive to ER stress (Figure 7A) (Chen & Brandizzi, 2012). This data suggests that the Gγ subunits are crucial for UPR. Taken together, these results suggest that the Gγ subunits inhibit the autophagic degradation of FLS2 at the ER membrane.

## DISCUSSION

The heterotrimeric G protein complex functions as a signal transducer at the plasma membrane in many signaling pathways including plant growth, development, and immunity. However, the downstream effectors of the heterotrimeric G proteins remains unknown. Moreover, the low number of heterotrimeric G proteins suggest that the current heterotrimeric G proteins may have additional functions outside of signal transduction at the plasma membrane or that there are additional undiscovered heterotrimeric G protein subunits (Miller et al., 2019). This study demonstrates that the Gγ subunits AGG1 and AGG2, which form an obligate heterodimer with the Gβ subunit AGB1, affect FLS2 protein levels under native growth conditions. Specifically, the *agg1 agg2* double-mutant has reduced FLS2 protein levels under normal growth conditions. However, the *agg1 agg2* mutant shows a decrease in immune signaling when treated with physiological levels of flagellin. While the loss of AGG1 and AGG2 debilitates immune signaling, the reduction of FLS2 protein levels contributes partially to the decrease in immune signaling, suggesting that the Gβγ heterodimer serves two functions in immunity: to transduce immune signals from FLS2 at the plasma membrane and maintain FLS2 protein levels.

The reduction of FLS2 protein levels in the *agg1 agg2* mutant is caused by increased FLS2 protein degradation. FLS2 is actively degraded by the proteasome and autophagy pathways in the *agg1 agg2* mutant. However, FLS2 levels can be rescued in *agg1 agg2* by inhibiting either the proteasome or autophagy pathway. Inhibiting both the proteasome and autophagy pathways increases FLS2 levels the most in the *agg1 agg2* mutant compared to inhibiting either the proteasome or autophagy alone. This suggests that in the absence of the Gβγ heterodimer, FLS2 is actively targeted by both protein degradation pathways. Inhibiting both the proteasome and autophagy pathways in the *agg1 agg2* mutant is also sufficient to rescue flagellin perception in this mutant, further supporting that the Gβγ heterodimer prevents proteasomal and autophagic degradation of FLS2. A recent report showed that the receptor-like cytoplasmic kinase (RLCK) BOTRYTIS-INDUCED KINASE 1 (BIK1) protein stability is dependent on the presence of the Gβγ heterodimer (Liang et al., 2016). In the absence of flagellin, BIK1 associates with FLS2 and its co-receptor BAK1 (Lu et al., 2010; Zhang et al., 2010). Upon flagellin perception, BAK1 phosphorylates BIK1 causing BIK1 to dissociate from the FLS2-BAK1 complex (Lu et al., 2010; Zhang et al., 2010). Together these findings support the hypothesis that the Gβγ heterodimer promotes the stability of membrane proteins such as FLS2 and BIK1.

The Gβγ heterodimer is crucial in preventing premature FLS2 protein degradation by the proteasome and autophagy. When either the *atg7* or *atg3* autophagy single mutants are crossed to the *agg1 agg2* mutant background, the autophagy machinery is disabled in the *agg1 agg2* mutant. FLS2 levels are rescued in these *agg1 agg2 atg7* and *agg1 agg2 atg3* triple mutants. Accordingly, MAPK activation upon flagellin treatment and resistance against *P. syringae* are also rescued in these triple mutants. The *agg1 agg2 atg7* and *agg1 agg2 atg3* triple mutants show that *AGG1* and *AGG2* are part of a signaling pathway that requires autophagy. Furthermore, the Gγ subunits localize and function at the endoplasmic reticulum membrane. AGG1 and AGG2 function in the unfolded protein response as the *agg1 agg2* mutant is hypersensitive to tunicamycin, which induces ER stress. This suggests that AGG1 and AGG2 function at the ER and inhibit the autophagic degradation of FLS2 protein. However, further research is need to determine the mechanism by which AGG1 and AGG2 prevent the proteasomal and autophagic degradation of FLS2.

We have shown that the Gβγ heterodimer functions to maintain plant sensitivity against flagellin by inhibiting the protein degradation of FLS2 under normal growth conditions. However, the mechanism by which the Gβγ heterodimer prevents FLS2 degradation remains elusive. We propose that the Gβγ heterodimer binds to the FLS2 kinase domain at the ER membrane to avoid inadvertent FLS2 activation while it travels through the secretory pathway to the plasma membrane. This model is supported by evidence demonstrating that the Gβγ heterodimer interacts directly with the FLS2 kinase domain at the plasma membrane along with the Gα subunit (Liang et al., 2016; Xu et al., 2017). Moreover, previous reports show that activation of FLS2 causes its subsequent degradation either through the proteasome or autophagic vesicles (Robatzek et al., 2006; Lu et al., 2011; Beck et al., 2012). Together, this suggests that the heterotrimeric G protein complex may bind to the FLS2 kinase domain at the ER membrane. In doing so, premature FLS2 activation is blocked by the heterotrimeric G protein complex, preventing the degradation of FLS2 by the proteasome and autophagy (Figure 8). As a result, FLS2 protein levels are maintained to ensure plant sensitivity against the PAMP, flagellin. Taken together, our results show that the plant canonical G proteins have additional functions in addition to signal transduction at the plasma membrane.

**Figure 8.**
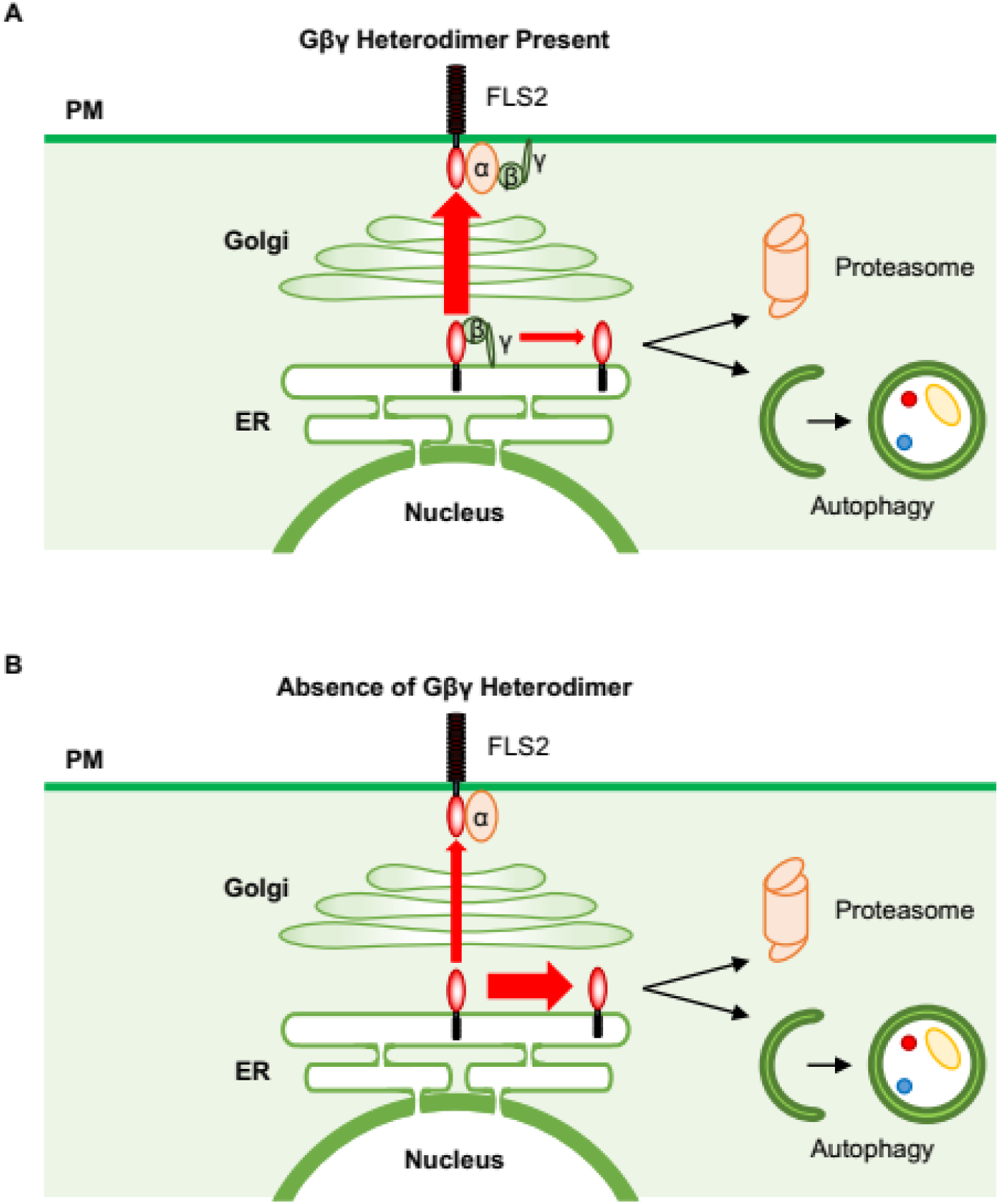
Model of how Gβγ heterodimer regulates FLS2 at the ER to prevent proteasomal and autophagic degradation of FLS2. (A) The Gβγ heterodimer binds the FLS2 kinase domain on the cytoplasmic side of the ER membrane. This inhibits FLS2 auto-activation and prevents FLS2 proteasomal and autophagic degradation as FLS2 travels from the ER to the plasma membrane. (B) In the absence of the Gβγ heterodimer, the FLS2 kinase domain auto-activates at the ER, resulting in FLS2 degradation by autophagy and the proteasome. This in turn causes a reduced amount of total FLS2 levels and contributes to the reduction of immune signaling.

## CONCLUSION

This study provides evidence that the Gβγ heterodimer maintains immunocompetency by regulating the protein degradation of the pattern recognition receptor FLS2 at the ER membrane. The Gβγ heterodimer may regulate FLS2 degradation by binding FLS2 at the ER membrane while FLS2 traverses through the secretory pathway to the plasma membrane. The binding of Gβγ heterodimer to the FLS2 kinase domain prevents premature FLS2 activation and its subsequent degradation by the proteasome and autophagy as the ectodomain of FLS2 matures through the secretory pathway. However, future studies are necessary to validate this hypothesis. One approach would be to create an *fls2 agg1 agg2* triple mutant and express a kinase dead FLS2_S938A_ (Cao et la., 2013) protein under the native *FLS2* promoter. FLS2 protein levels should be measured in this line under normal growth conditions to see if there is a change in FLS2 levels. Furthermore, more research is needed to determine if the Gβγ heterodimer is able to regulate other pattern recognition receptors in a similar manner to FLS2.

As the heterotrimeric G proteins have only ever been thought to function at the plasma membrane, we have shed light on the capabilities of the plant heterotrimeric G proteins in immunity that go beyond its function at the plasma membrane in signal transduction. This work shows the expanded capacity of the heterotrimeric G proteins from signal transduction at the plasma membrane to a novel function preventing the protein degradation of FLS2 at the ER membrane.

## MATERIALS & METHODS

### Plant materials and growth conditions

Surface-sterilized seeds of *Arabidopsis thaliana* accession Columbia-0 (Col-0) were stratified for at least 2 days and sown in 12- or 24-well microtiter plates sealed with parafilm. Each well of the 12- or 24-well plate contained 12 and 5 seeds, respectively, with 1 and 0.5 mL of filter-sterilized 0.5X MS liquid (pH 5.7–5.8) [4.43 g/L Murashige and Skoog basal medium with vitamins (Murashige and Skoog, 1962) (Phytotechnology Laboratories, Shawnee Missions, KS), 0.05% (w/v) MES hydrate, 0.5% (w/v) sucrose], respectively. Alternatively, surface-sterilized and stratified seeds were sown on MS agar plates [0.5X MS, 0.75% (w/v) agar (PlantMedia, Chiang Mai, Thailand)] and sealed with parafilm. Unless otherwise stated, sample-containing plates were placed on grid-like shelves over water trays on a Floralight cart (Toronto, Canada), and plants were grown at 21°C and 60% humidity under a 12-hr light cycle (70–80 μE m^−2^ s^−1^ light intensity). Unless otherwise stated, media in microtiter plates were exchanged for fresh media on day 7. For bacterial infection experiments, *Arabidopsis* plants were grown on soil [3:1 mix of Fafard Growing Mix 2 (Sun Gro Horticulture, Vancouver, Canada) to D3 fine vermiculite (Scotts, Marysville, OH)] at 22°C daytime/18°C nighttime with 60% humidity under a 12-hr light cycle (100 µE m^−2^ s^−1^ light intensity). *Nicotiana benthamiana* plants were grown on soil [3:1 mix] on a Floralight cart at 22°C under a 12-hr light cycle (100 µE m^−2^ s^−1^ light intensity) for 4 weeks.

The following Col-0 T-DNA insertion lines and mutants were obtained from the Arabidopsis Biological Resource Center (ABRC, Columbus, Ohio): *agb1-1* (CS3976)*, agb1-2* (CS6535), *agg1-1c* (CS16550), *agg2* (SALK_039423), *agg1-1c/agg2-1* (CS16551), *atg7* (SAIL_11_H07), *fls2* (SAIL_691_C4). The *atg3-1* mutant was isolated by Farmer et al., 2013.

### Vector construction and transformation

To generate estradiol-inducible C-terminally tagged *GFP* and *RFP* (*XVE:X-G/RFP*) and *35S:YFP-AGB1* DNA constructs, *attB* sites were added via PCR-mediated ligation to the coding sequences of cDNAs, and the modified cDNAs were recombined into pDONR221 entry vector and then into pABindGFP, pABindRFP (Bleckmann et al., 2010) and pB7WGY2 (Karimi et al., 2002) destination vectors, according to manufacturer’s instructions (Gateway manual; Invitrogen, Carlsbad, CA). *XVE:AGG1-RFP, XVE:FLS2-RFP,* and *35S:YFP-AGB1* constructs were introduced into *agg1-1c agg2-1* or *agb1-2* plants via *Agrobacterium*-mediated floral dip method (Clough and Bent, 1998), and transformants were selected on agar media containing 15 µg/mL hygromycin B (Invitrogen) or 15 µg/mL glufosinate (Cayman Chemical, Ann Arbor, MI). Transgene expression was induced 48 hr (or 5-6 days for growth assays) after elicitation with 20 μΜ β-estradiol (2 mM stock solution in DMSO; Sigma-Aldrich, St. Louis, MO). Transient expression of *XVE:X-G/RFP* constructs in *Nicotiana benthamiana* leaves was performed as previously described (Bleckman et al., 2010) with the following modification: transformed *Agrobacterium* strains were grown in LB medium supplemented with 50 µg/mL rifampicin, 30 µg/mL gentamycin, kanamycin 50 μg/mL and 100 µg/mL spectinomycin, in the absence of a silencing suppressor, to an OD_600_ of 0.7. Transgene expression was induced 10 hr (for co-immunoprecipitation) and 4-8 hr (for microscopy) after spraying with 20 µM β-estradiol and 0.1% Tween-20.

### Flg22-induced seedling growth inhibition

Three-day-old seedlings in 24-well microtiter plates were elicited with water or 100 nM flg22 (QRLSTGSRINSAKDDAAGLQIA; Genscript, Nanjing, China) for 6 days. Fresh weights were measured from 9-day-old seedlings that were dried between paper towels for a few seconds.

### Flg22-induced MAPK activation

9-day-old seedlings were elicited with 100 nM flg22 for 5, 15, and/or 30 min. MAPK activation assay was performed as previously described (Lawerence et al., 2017). 20 μl of supernatant was loaded onto a 10% SDS-PAGE gel, and the separated proteins were transferred to PVDF membrane (Millipore) and probed with phosphor-p44/p42 MAPK (Cell Signaling Technology, Danvers, MA) and MPK3 antibodies (Sigma-Aldrich, St. Louis, MO) at 1:2000 dilution in 5% (w/v) nonfat milk in 1X PBS. The combined signal intensities of phosphorylated MPK3/4/6 were quantified using NIH ImageJ and normalized to that of total MPK3 (loading control).

### Flg22-induced Callose Deposition

9-day-old seedlings were elicited with 1 μM flg22 for 16-18 hr. Callose deposition staining was performed as previously described (Clay et al., 2009). Callose deposits were viewed on a Zeiss (Oberkochen, Germany) AxioObserver D1 fluorescence microscope under UV illumination with Filter Set 49 (excitation filter 365 nm; dichroic mirror 395 nm; emission filter 445/50 nm). Callose deposits were quantified using NIH ImageJ.

### Total protein extraction, SDS-PAGE, and western blotting

Total protein was extracted from snap-frozen seedlings into 80 µL of extraction buffer [50 mM Tris-Cl (pH 7.5), 50 mM DTT, 4% (w/v) SDS, 10% (v/v) glycerol] using a 5-mm stainless steel bead and ball mill (20 Hz for 3 min). Samples were centrifuged briefly, incubated at 95°C for 10 min, and centrifuged at 12,000 x g for 8 min to precipitate insoluble material. 5 or 10 μL of extract were loaded onto a 8.5% SDS-PAGE gel, and the separated proteins were transferred to PVDF membrane (Millipore, Billerica, MA), stained with Ponceau S for labeling of total protein (loading control), and probed with FLS2 (antigen: CTKQRPTSLNDEDSQ; Genscript) at a 1:1000 (FLS2) dilution in 5% (w/v) nonfat milk in 1X PBS. Signal intensities of immuno-detected proteins were quantified using NIH ImageJ and normalized to that of the loading control.

### RNA isolation and quantitative PCR (qPCR)

Total RNA was extracted into 1 mL of TRIzol reagent (Invitrogen) according to manufacturer’s instructions. 2 µg of total RNA was reverse-transcribed with 3.75 µM random hexamers (Qiagen, Hilden, Germany) and 20 U of ProtoScript II (New England Biolabs, Boston, MA). The resulting cDNA:RNA hybrids were treated with 10 U of DNase I (Roche) for 30 min at 37°C, and purified on PCR clean-up columns (Macherey-Nagel, Düren, Germany). qPCR was performed with Kapa SYBR Fast qPCR master mix (Kapa Biosystems, Wilmington, MA) and CFX96 or CFX384 real-time PCR machine (Bio-Rad, Hercules, CA). The thermal cycling program is as follows: 95°C for 3 min; 45 cycles of 95°C for 15 sec and 53°C or 55°C for 30 sec; a cycle of 95°C for 1 min, 53°C for 1 min, and 70°C for 10 sec; and 50 cycles of 0.5°C increments for 10 sec. Biological replicates of control and experimental samples, and three technical replicates per biological replicate were performed on the same 96- or 384-well PCR plate. Averages of the three Ct values per biological replicate were converted to differences in Ct values relative to that of control sample. The Pfaffl method (Pfaffl, 2001) and calculated primer efficiencies were used to determine the relative fold increase of the target gene transcript over the housekeeping *eIF4AI* gene transcript for each biological replicate. Primer sequences and efficiencies are listed in Table 1.

**Table 1.**
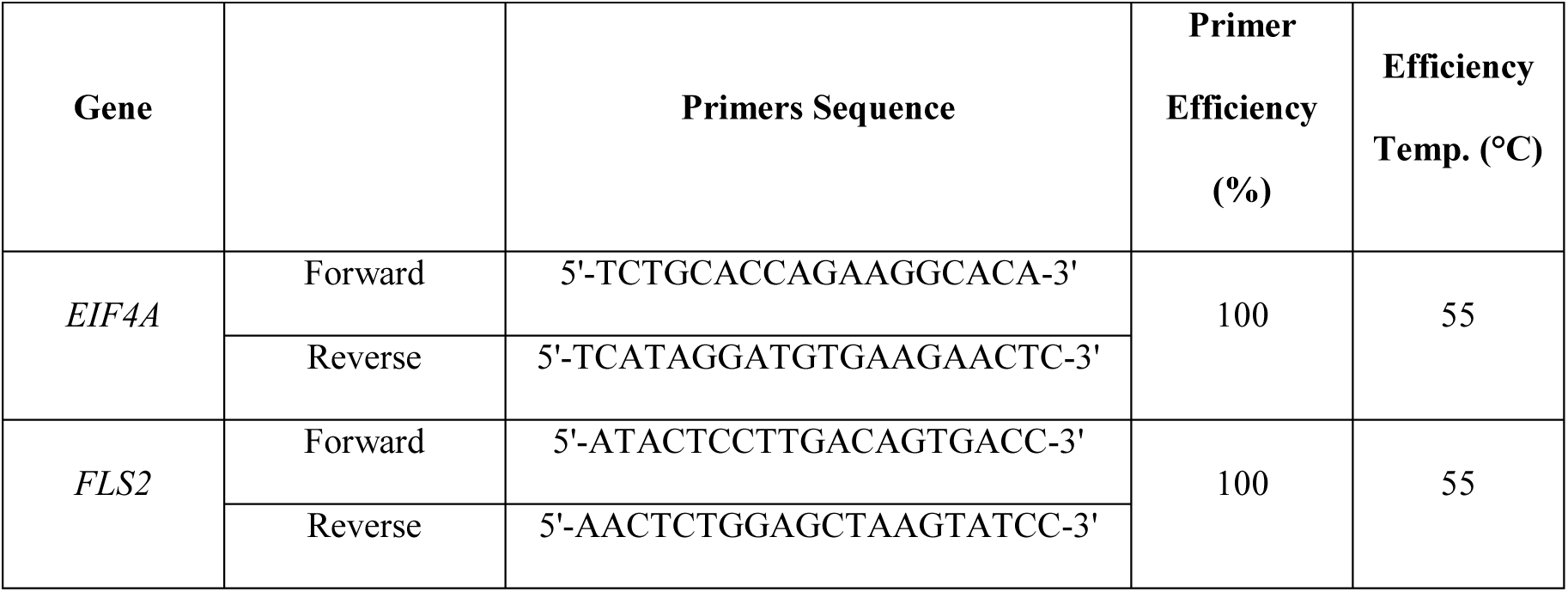
Q-PCR primer sequences and efficiencies.

### Inhibition of protein translation and degradation

For FLS2 western blots, 9-day-old seedlings were treated with 50 µM cycloheximide for 0, 2, and 16 hr. 9-day-old seedlings were also treated with DMSO (mock), 50 µM MG132 (50 mM stock solution in DMSO; Selleck Chemicals, Houston, TX), 20 µM E-64d (20 mΜ stock solution in DMSO; Cayman Chemical), and/or 2 µM concanamycin A (200 μM stock solution in DMSO; Santa Cruz Biotechnology) for 24 hr. For growth inhibition, 3-day-old seedlings were elicited with 100 nM flg22 or water for 2 days, and then treated with DMSO, 50 nM MG132, and/or 20 nM E-64d for 4 days.

### Bacterial infection

*Pseudomonas syringae* pv *tomato* DC3000 (*Pto* DC3000) was used for bacterial infections. *Pto* DC3000 was grown in Luria-Bertani medium and 25 µg/mL rifampicin (Sigma-Aldrich) overnight, washed in sterile water twice, and re-suspended in water to an OD_600_ of 0.002. Adult leaves of 4-5-week-old plants were surface-inoculated with the bacterial inoculum (OD_600_ = 0.002 or 10^6^ colony-forming units [CFU]/cm^2^ leaf area) in the presence of 0.0075% Silwet L-77 (Phytotechnology Laboratories) for 15 s and incubated in 0.8% (w/v) tissue culture agar plates for 3 to 4 d. Infected leaves were surface-sterilized in 70% ethanol for 10 s, washed in sterile water, and dried on paper towels. The bacteria were extracted into water using an 8-mm stainless steel bead and a ball mill (25 Hz for 3 min). Serial dilutions of the extracted bacteria were plated on Luria-Bertani agar plates for CFU counting

### Confocal microscopy

Live epidermal root cells of 5-day-old *Arabidopsis* seedlings and 4-week-old *N. benthamiana* leaves were imaged using a 40X 1.0 numerical aperture Zeiss water-immersion objective and a Zeiss LSM 510 Meta confocal microscopy system. GFP and RFP were excited with a 488-nm argon laser and 561-nm laser diode, respectively. GFP and RFP emissions were detected using a 500-550 nm and 575-630 nm filter sets, respectively. Plasmolysis was induced by 5-10 min treatment of *N. benthamiana* leaf strips with 0.8 M mannitol, and co-localization of GFP/RFP-tagged proteins to Hechtian strands was made visible by over-exposing confocal images using ZEN software.

### ER stress induction

Seedlings were grown on MS agar supplemented with 0, 25, 50, and 100 ng/mL tunicamycin (0.5 mg/mL stock solution in DMSO; Sigma-Aldrich) for 14 days.

## ACKNOWLEDGEMENTS

We thank Nozomu Koizumi for the *ire1a-2/ire1b-2* (SALK_018112, GABI_638B07) mutant and Bonnie Bartel for the *atg3-1* mutant. We thank Simon Rüdiger for the pABindGFP and pABindmCherry vectors. We thank Josh Gendron for help with interpreting the data. The work was supported by T32 GM007499 (to S.A.L) and T32 GM007223 (to J.C.M)

---

**Supplementary Figure S1.**
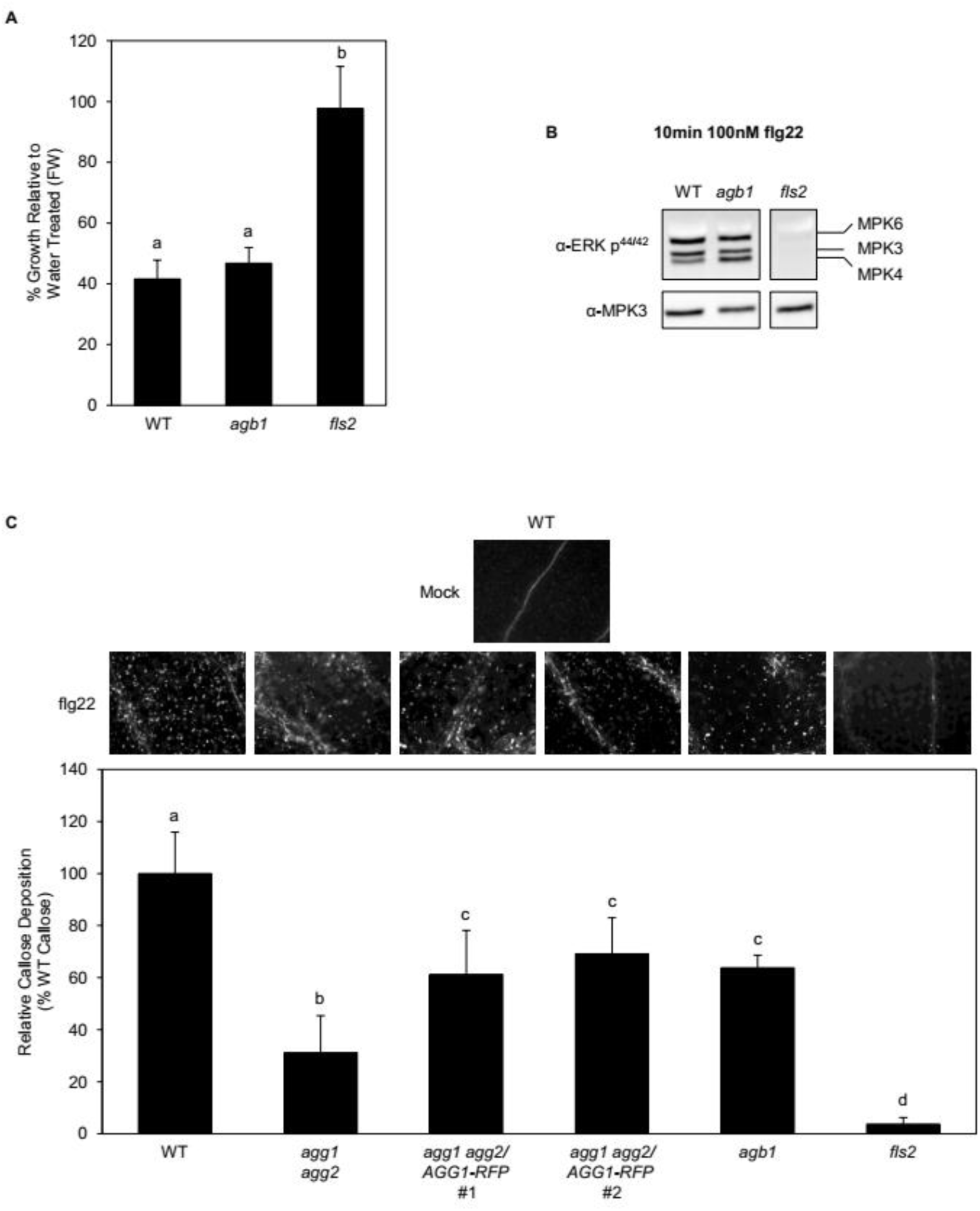
The Gβγ heterodimer is crucial in immune signaling at physiological flagellin levels. Growth inhibition analysis of 9-day-old seedlings pre-treated with water (control) or 100 nM flg22 for 6 days. Data represent mean ± SD of four (*fls2*) or five (all others) replicates of five seedlings. (B) Immunoblot analysis of phosphorylated, active MAPKs in 9-day-old WT*, agb1*, and *fls2* in response to 100 nM flg22 for 10 minutes. Total MPK3 was used as a loading control. (C) Callose deposition analysis of 9-day-old WT, *agg1 agg2*, *agg1 agg2/XVE:AGG1-RFP*, *agb1,* and *fls2* plants in response to water (control) or 1 µM flg22 for 16-18 hr. Plants on the left side of the graph were pretreated with 20 µM β-estradiol for 48 hr. Data represent mean ± SE of 25 replicates. Different letters indicate significant differences (*P*-value <0.05, two-tailed *t* test).

**Supplementary Figure S2.**
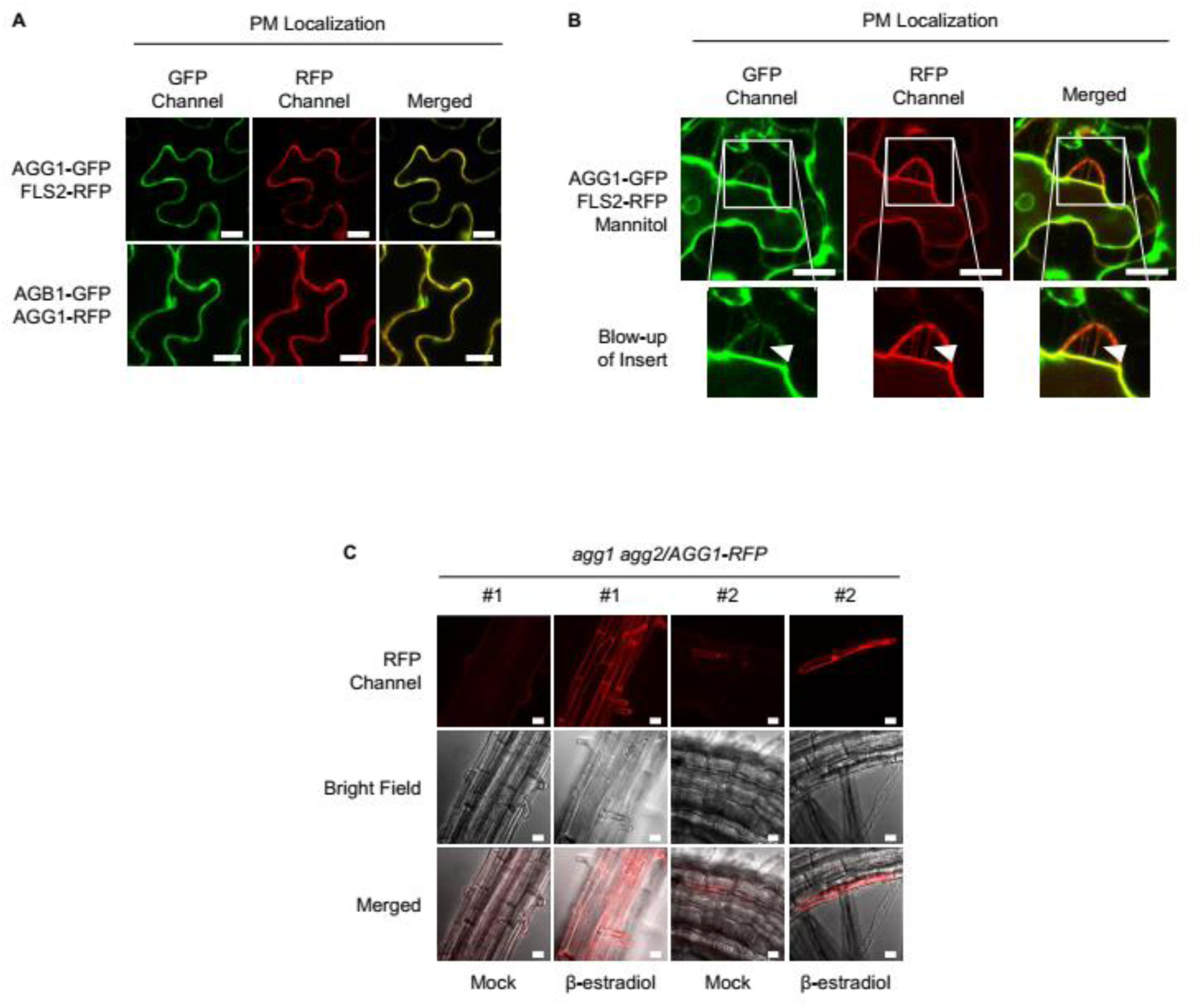
C-terminally tagged AGG1 does not affect the localization of AGG1. (A) Co-localization of AGG1 with FLS2 and AGB1 at the PM. (*B*) Co-localization of AGG1 with FLS2 at plasmolysis-induced Hechtian strands (indicated by white arrowheads). (*C*) Co-localization of AGG1-RFP to reticulate structures in 5-day-old *agg1 agg2/XVE:AGG1-RFP* Arabidopsis plants pretreated with 20 μM β-estradiol for 48 hr. Experiments in (*A and B*) were performed on transfected *N. benthamiana* leaves pretreated with 20 µM β-estradiol for 4 hr. White bars in (*A–C*) represent 20 μm.

